# Resiniferatoxin (RTX) ameliorates acute respiratory distress syndrome (ARDS) in a rodent model of lung injury

**DOI:** 10.1101/2020.09.14.296731

**Authors:** Taija M. Hahka, Zhiqiu Xia, Juan Hong, Oliver Kitzerow, Alexis Nahama, Irving H. Zucker, Hanjun Wang

## Abstract

Acute lung injury (ALI) is associated with cytokine release, pulmonary edema and in the longer term, fibrosis. A severe cytokine storm and pulmonary pathology can cause respiratory failure due to acute respiratory distress syndrome (ARDS), which is one of the major causes of mortality associated with ALI. In this study, we aimed to determine a novel neural component through cardiopulmonary spinal afferents that mediates lung pathology during ALI/ARDS. We ablated cardiopulmonary spinal afferents through either epidural T1-T4 dorsal root ganglia (DRG) application or intra-stellate ganglia delivery of a selective afferent neurotoxin, resiniferatoxin (RTX) in rats 3 days post bleomycin-induced lung injury. Our data showed that both epidural and intra-stellate ganglia injection of RTX significantly reduced plasma extravasation and reduced the level of lung pro-inflammatory cytokines providing proof of principle that cardiopulmonary spinal afferents are involved in lung pathology post ALI. Considering the translational potential of stellate ganglia delivery of RTX, we further examined the effects of stellate RTX on blood gas exchange and lung edema in the ALI rat model. Our data suggest that intra-stellate ganglia injection of RTX improved pO_2_ and blood acidosis 7 days post ALI. It also reduced wet lung weight in bleomycin treated rats, indicating a reduction in lung edema. Taken together, this study suggests that cardiopulmonary spinal afferents play a critical role in lung inflammation and edema post ALI. This study shows the translational potential for ganglionic administration of RTX in ARDS.

## Introduction

Respiratory failure due to acute respiratory distress syndrome (ARDS) is one of the major causes of mortality associated with acute lung injury (ALI) including COVID-19.^1-5^ Most forms of ALI/ARDS are also associated with acute cytokine release, pulmonary edema and in the longer term, fibrosis.^5,6^ However, the mechanisms underlying these pathological changes in the lungs during ALI/ARDS are not fully understood. In particular, a neural component that mediates lung pathology during ALI/ARDS has been less considered.

Sensory neurons innervating the heart and lung enter the central nervous system by one of two routes; through the vagus nerve into the brain stem (medulla) with cell bodies residing in the nodose ganglia and directly into the spinal cord where cell bodies reside in the Dorsal Root Ganglia (DRG). Afferents are composed of elements that respond to a variety of sensory modalities including mechanical deformation, heat, cold, pH, and inflammatory mediators, just to name a few.^7^ The reflex effects following stimulation of these afferents depends on the type of stimulus and the neural pathway involved. Activation of vagal afferent pathways tends to be sympatho-inhibitory and anti-inflammatory.^8, 9^ On the other hand, activation of spinal afferents tends to be sympatho-excitatory and pro-inflammatory.^10-15^ It is well known that small diameter spinal Transient Receptor Vanilloid 1 (TRPV1)-positive afferent c-fibers contain neuropeptides such as substance P (SP) and calcitonin gene related peptide (CGRP).^16^ These peptides tend to dilate adjacent vasculature and increase microvascular permeability.^16^ In the lung, this can cause pulmonary edema resulting in reduced oxygen diffusion and promote immune cell infiltration resulting in neural inflammation. Therefore, in the current study we hypothesized that ablation of lung afferent innervation (thoracic spinal) by application of an ultrapotent, selective afferent neurotoxin, resiniferatoxin (RTX) will modify the course of the pathology including lung edema and local pulmonary inflammation associated with progressive ALI.

## Methods

All animal experimentation was approved by the Institutional Animal Care and Use Committee of the University of Nebraska Medical Center and performed in accordance with the National Institutes of Health’s *Guide for Use and Care of Laboratory Animals* and with ARRIVE guidelines.^17, 18^ Experiments were performed on adult, male, 200-250g Sprague-Dawley rats purchased from the Charles River Laboratories. Animals were housed on-site and given a one-week acclimation period prior to experimentation. Food and water were supplied *ad libitum*, and rats were on 12-hour light/dark cycles.

### Rat model of lung injury

Rats were randomized into three groups and evaluated at 1-week post-instillation as follows: sham rats, bleomycin (Bleo)-exposed rats with saline (epidural or intra-stellate injection), and Bleo-exposed rats with RTX (epidural or intra-stellate injection). Bleo (2.5 mg/kg, ∼0.15 mL) was instilled intra-tracheally to the lungs under 3% isoflurane anesthesia. Sham control rats underwent intra-tracheal instillation of saline. Animals were treated with RTX or vehicle (phosphate buffered saline) by either the epidural T1-T4 DRGs route (6 µg/ml, 10µl/per ganglia) or intra-stellate ganglia administration (50 µg/ml, 5 µl/per side) 3 days following bleomycin delivery (Figure 1).

**Figure 1.**
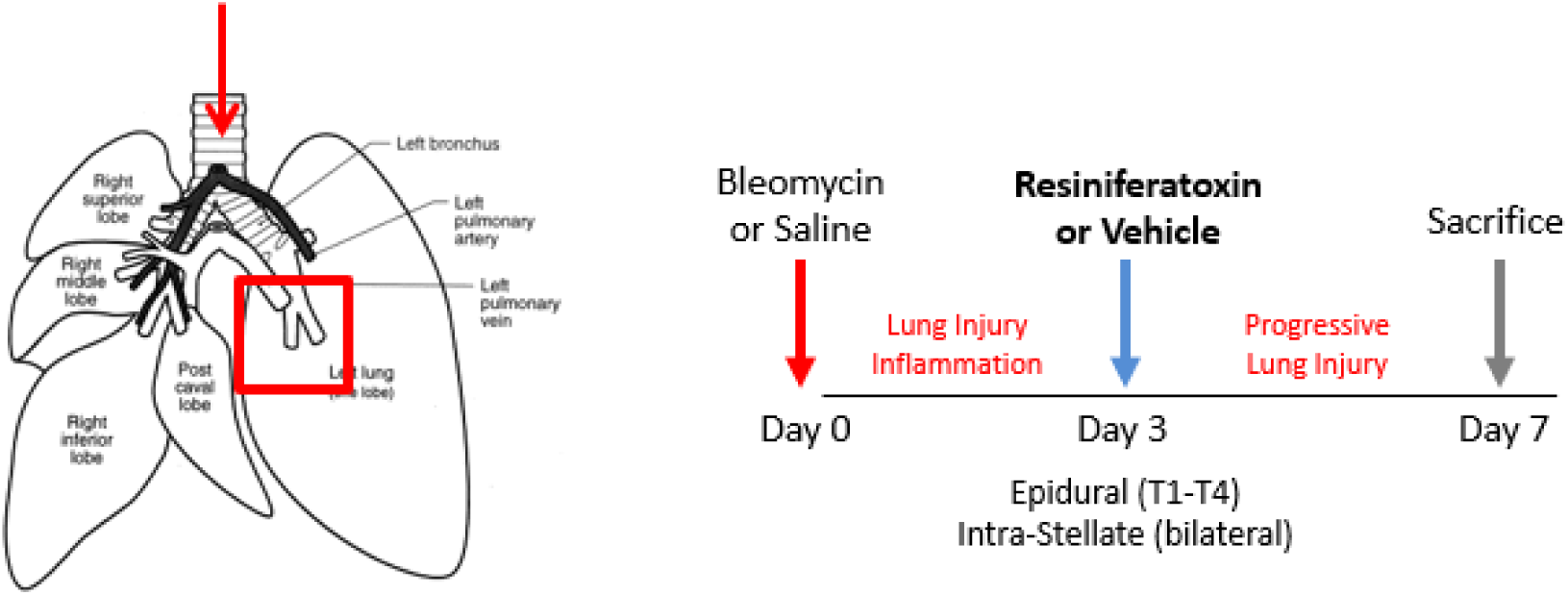
Schematic diagram of study design. Left panel: red arrow indicates that bleomycin (2.5 mg/kg, ∼0.15 mL) was administrated intra-tracheally to the lungs. Red square shows the location where lung tissue was collected for cytokine measurement. Right panel: At day 0, bleomycin or saline was given intra-tracheally; at day 3, resiniferatoxin or vehicle was given into epidural space or into stellate ganglia; at say 7, the rats were sacrificed.

### Epidural application of RTX

In a pilot experiment, the upper thoracic spinal afferents were ablated by epidural application of RTX as previously described.^11^ Briefly, rats were anesthetized using 2%-3% isoflurane:oxygen mixture. Rats were placed in the prone position and a small midline incision was made in the region of the T13-L1 thoracic vertebrae. Following dissection of the superficial muscles, two small holes (approximately 2 mm x 2 mm) were made in the left and right sides of T13 vertebrae. A polyethylene catheter (PE-10) was inserted into the subarachnoid space via one hole and gently advanced about 4cm approximating the T1 level. The upper thoracic sympathetic afferent ganglia were ablated by injecting resiniferatoxin (RTX; Sigma Aldrich), an ultra-potent agonist of the TRPV1 receptor into the aubarachnoid space via the catheter. RTX (1 mg; Sigma Aldrich) was dissolved in a 1:1:8 mixutre of ethanol, Tween 80 (Sigma-Aldrich), and isotonic saline. The first injection of RTX (6 µg/ml, 10ul) was made at a very slow speed (∼ 1 minute) to minimize the diffusion of the drug. The catheter was then pulled back to T2, T3 and T4, respectively to perform serial injections (10 ul/each) at each segment. The catheter was withdrawn and the same injections were repeated on the other side. Silicone gel was used to seal the hole in the T13 vertebra. The skin overlying the muscle were closed with a 3-0 polypropylene simple interrupted suture, and betadine was applied to the wound. For post-procedure pain management, buprenorphine (0.05 mg/kg) was subcutaneously injected immediately after surgery and twice daily for 2 days.

### Intra-stellate injection of RTX

Rats were anesthetized using 2%-3% isoflurane:oxygen mixture. After the trachea was cannulated mechanical ventilation was started (model 683, Harvard Apparatus, South Natick, MA). The skin from the rostral end of the sternum to the level of third rib was incised. Portions of the superficial and deep pectoral muscles and the first intercostal muscles were cut and dissected. To localize the left or right stellate ganglion, the left or right precava vein were separated with a hooked glass or steel rod laterally away from the brachiocephalic artery to expose the internal thoracic artery and the costocervical artery, which are descending branches of the right subclavian artery. Stellate ganglia and ansa subclavia are located medially to the origins of the internal thoracic and costocervical arteries. Then, RTX (5µl, 50 mg/ml) was injected into the ganglia with a 5-µl Hamilton syringe (Microliter #95, Hamilton, Reno, NV, USA.) over 30 s bilaterally. An image of this procedure is shown in **Figure 2**. Following these maneuvers, the thorax between the 1^st^ and 2nd intercostal spaces was closed with continuous 4-0 Dexon II coated braided absorbable polyglycolic acid suture and the skin was closed with 3-0 polypropylene suture and the chest evacuated. Betadine was applied to the wound and the rats were allowed to recover from the anesthesia. For post-procedure pain management, buprenorphine (0.05 mg/kg) was injected subcutaneously immediately after surgery and twice daily for 2 days.

**Figure 2.**
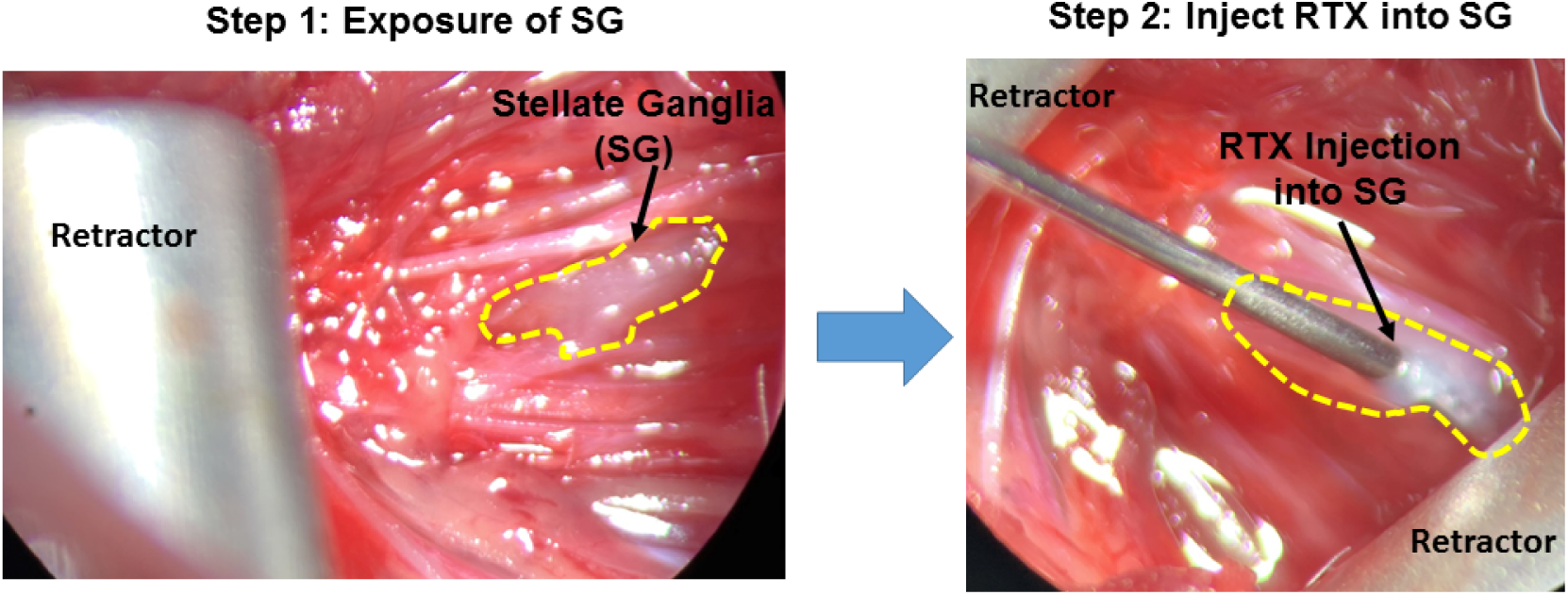
Procedure for stellate isolation and administration of vehicle or RTX. Step 1: stellate ganglia exposure. Black arrow shows that stellate ganglion is located medially to the origins of internal thoracic and costocervical arteries. Step 2: RTX injection. RTX (5µl, 50 mg/ml) was injected into the left and right stellate ganglions. Black arrow shows the tip of a 5-µl syringe inside the stellate ganglion.

### Blood gas analysis

The artery on the ventral aspect of the rat tail was used for the collection of small amounts of blood (∼0.1 mL) for analyzing arterial blood gas at day 7 post bleomycin. The animal was restrained with a commercial restrainer so that its tail was accessible. The tail was prepared aseptically by alternating alcohol prep pads and iodine prep pads three times and the artery was punctured using a 24 G needle. A small volume of blood (∼0.1 mL) was gently aspirated into the syringe for blood gas analysis (iSTAT, Abbott, Chicago, IL, USA). After sample collection, the needle was removed, and a gauze swab was pressed firmly on the puncture site to stop bleeding.

### Cytokine Assays

Lung and plasma cytokines were measured with R&D cytokine ELISA assays (Minneapolis, MN, USA) according to the manufacturer’s instructions. Organ weights were evaluated post-mortem.

### Lung plasma extravasation, tissue extraction and quantification of Evans blue

Rats were anaesthetized with pentobarbitone (40 mg/kg). Evans Blue, 20 mg/kg (10 mg/ml, dissolved in saline + 100 IE per ml heparin) was administered intravenously. After 10 min, rats were euthanized by transcardial perfusion with PBS (0.01 M, pH 7.4). The lung was taken and first photographed. Then, lung samples were immediately weighed, placed in 2 ml of N,N’-dimethyl formamide and was cut into small pieces at 50 °C water bath overnight. The lung tissues were then centrifuged (1 min, 14,000 rpm) and the Evans blue content of the lungs in the supernatant was determined in a 96-well microplate reader (infinite M200, TECAN, Männedorf, CH, Switzerland) at 620 nm (100 µl sample per well). Extravasation of Evans blue was expressed as mg Evans blue/g of lung tissue, by comparing the experimental values with a known standard.

### Statistics

Statistical evaluation was analyzed using GraphPad Prism (GraphPad Software, San Diego, CA. Version 8). Differences between treatments were determined using a Mixed-effects model for repeated-measures ANOVA. For comparison between three groups (Sham, Bleo+Veh and Bleo+RTX experiments) both Tukey and Bonferroni corrections for multiple comparisons were used.

The procedure by which the stellate ganglia were exposed and injected in rats under anesthesia is shown in **Figure 2**.

## Results

Plasma extravasation (Evans Blue) was used to assess vascular permeability after acute lung injury. As shown in **Figure 3**, bleomycin-treated lungs exhibited a wide distribution of Evans blue areas in both sides. The highest intensity of Evans blue was shown at the medial aspect of each lung. The Evans blue areas were largely reduced following epidural RTX treatment at the 7-day time point after bleomycin administration (**Figure 3**).

**Figure 3.**
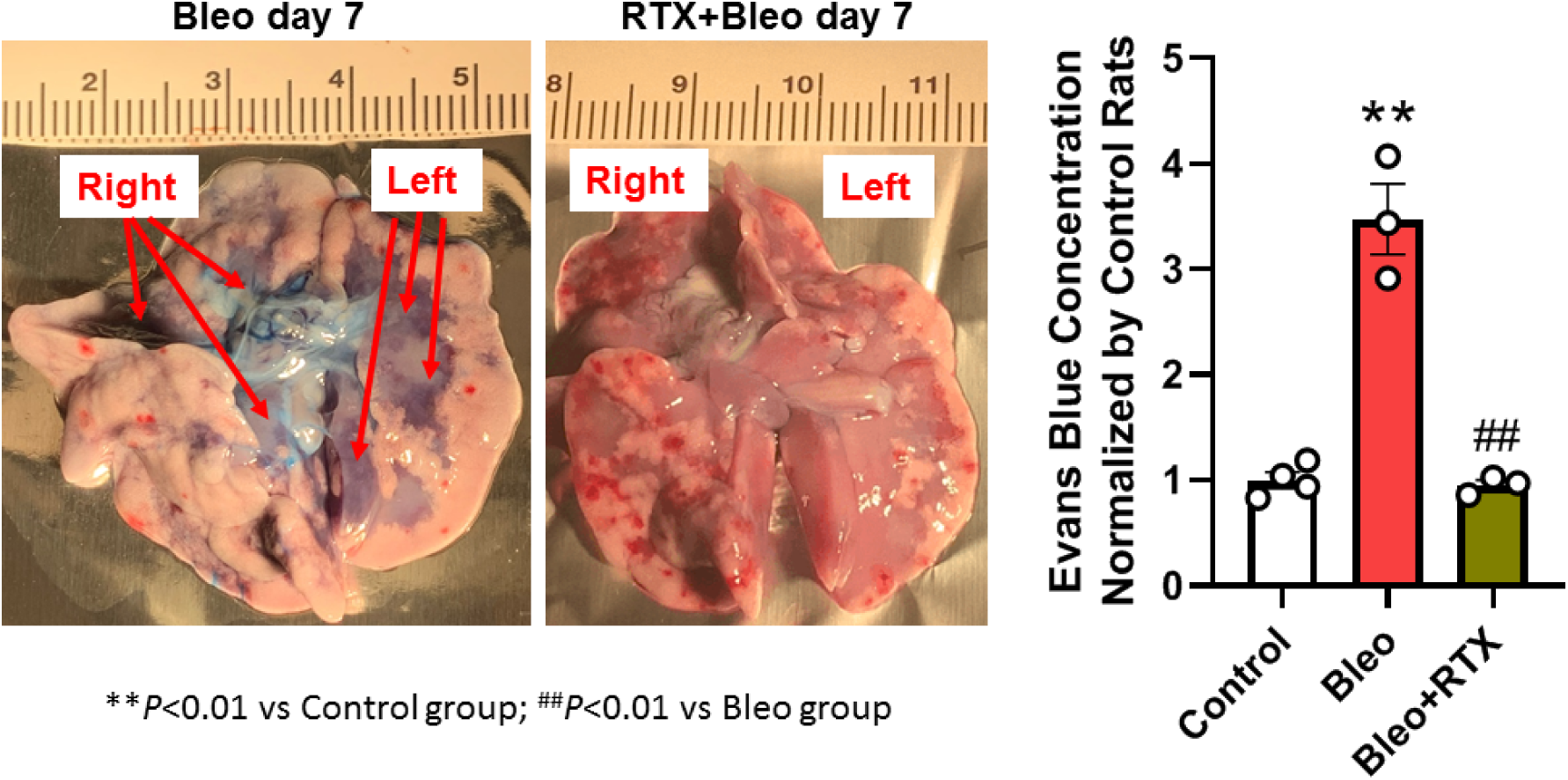
Plasma extravasation was reduced following epidural RTX treatment at the 7-day time point after bleomycin administration. Left panel shows representative images of the lungs from Bleo group and Bleo+RTX group. Right panel shows Evans blue concentration from Control, Bleo and Bleo+RTX rats. \*\**P*<0.01 vs. Control. ##*P*<0.01 vs. Bleo.

Three pro-inflammatory tissue cytokines were prevalent in the lung following Bleo treatment are shown in **Figure 4**. IL-6, IL-1ß and IFNý were markedly elevated following Bleo treatment. These cytokine levels were normalized in epidural RTX treated rats.

**Figure 4.**
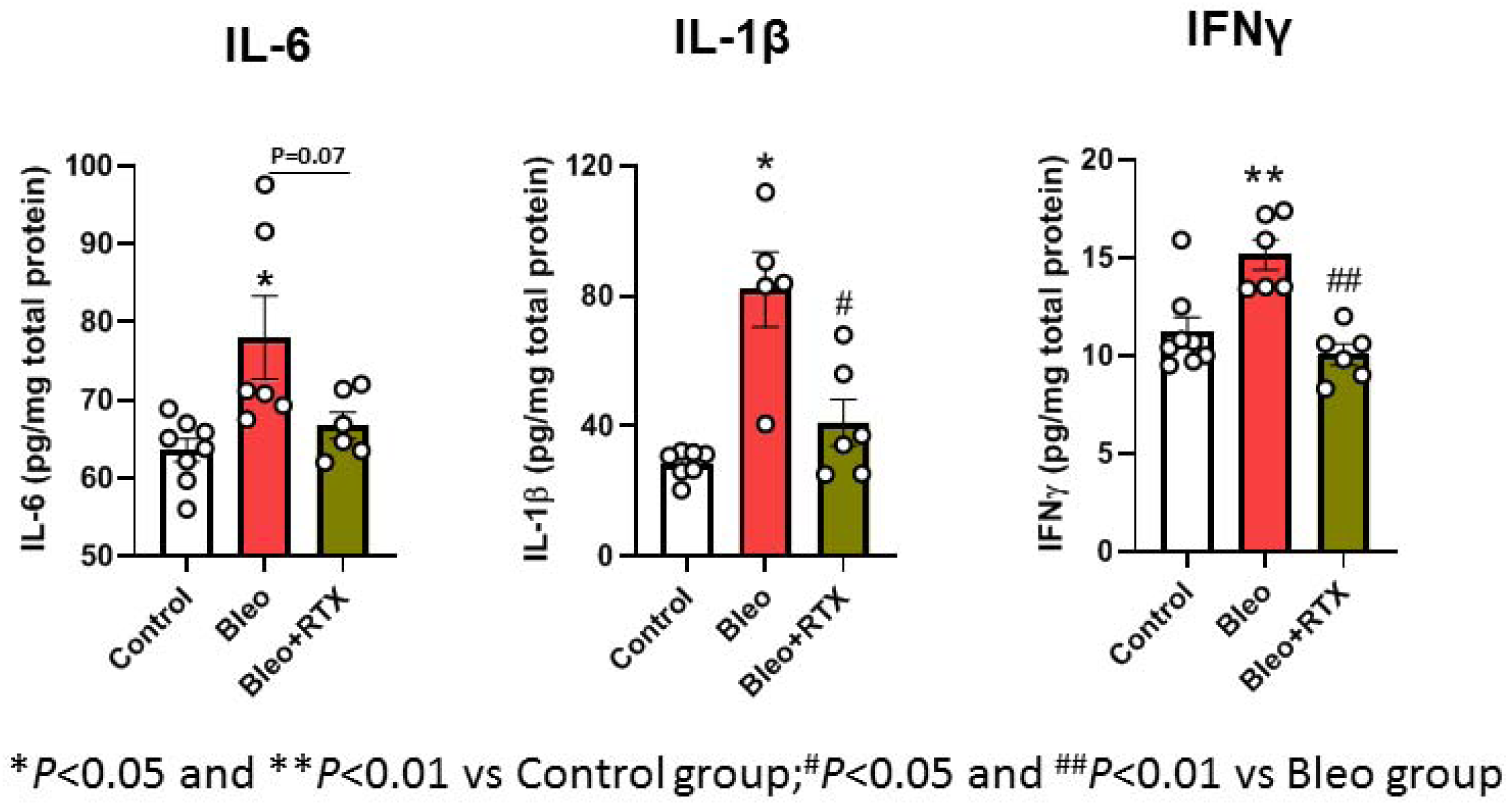
Day 7 lung tissue cytokine levels following day 3 vehicle or epidural RTX administration. IL-6, interleukin 6; IL-1β, interleukin 1 β; IFNγ, interferon γ. \**P*<0.05 and \*\**P*<0.01 vs. Control. ^#^*P*<0.05 and ^##^*P*<0.01 vs. Bleo.

Plasma extravasation in response to Bleo was also reduced after stellate injection of RTX (**Figure 5**). As can be seen, there was a marked reduction in Evans Blue dye in the lung following stellate injection of RTX.

**Figure 5.**
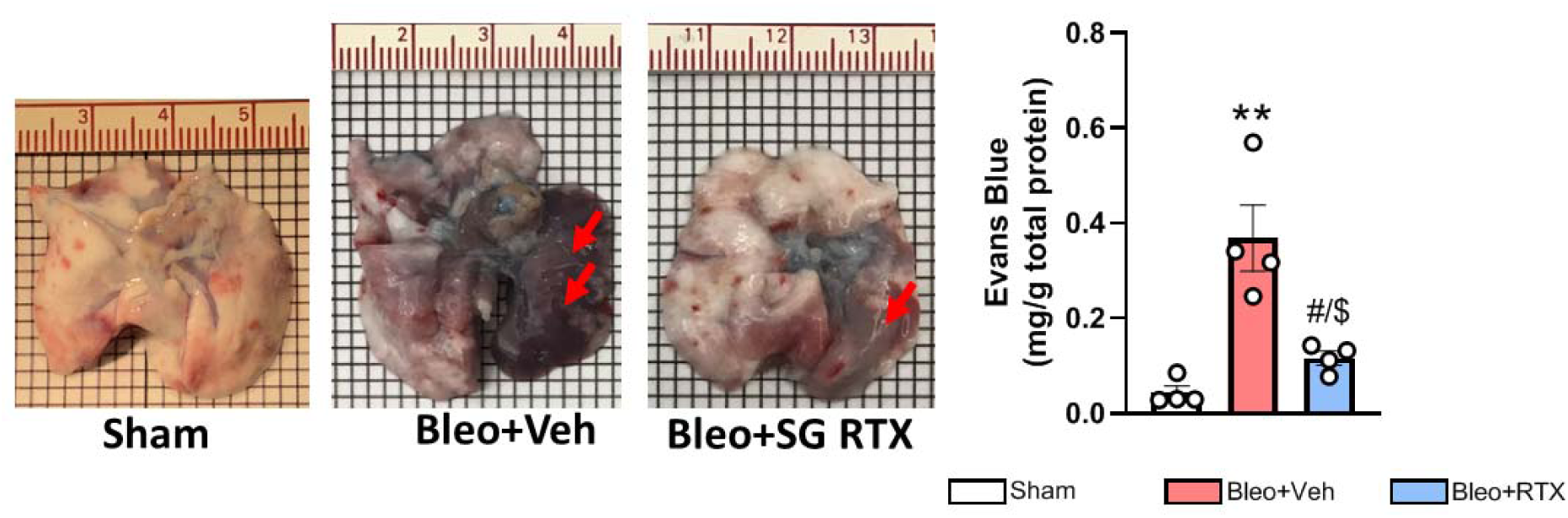
Evans blue extravasation was reduced following stellate RTX injection at the 7-day time point after Bleo administration. Left panel shows representative images of the lungs from Sham, Bleo+Veh group, and Bleo+RTX group. Red arrows point to areas of Evans blue extravasation. Right panel shows mean Evans blue concentration from each group. \*\**P*<0.01 vs. Sham. ^#^*P*<0.05 vs. Bleo+Veh. ^$^*P*<0.05 vs. Sham.

Arterial blood gas data were evaluated in rats treated with vehicle vs RTX intra-stellate. **Figure 6** shows that pCO_2_ was elevated and pO_2_ was reduced as was sO_2_ in Bleo vehicle treated rats. Stellate administration of RTX reversed these changes suggesting improved pulmonary function and gas exchange.

**Figure 6.**
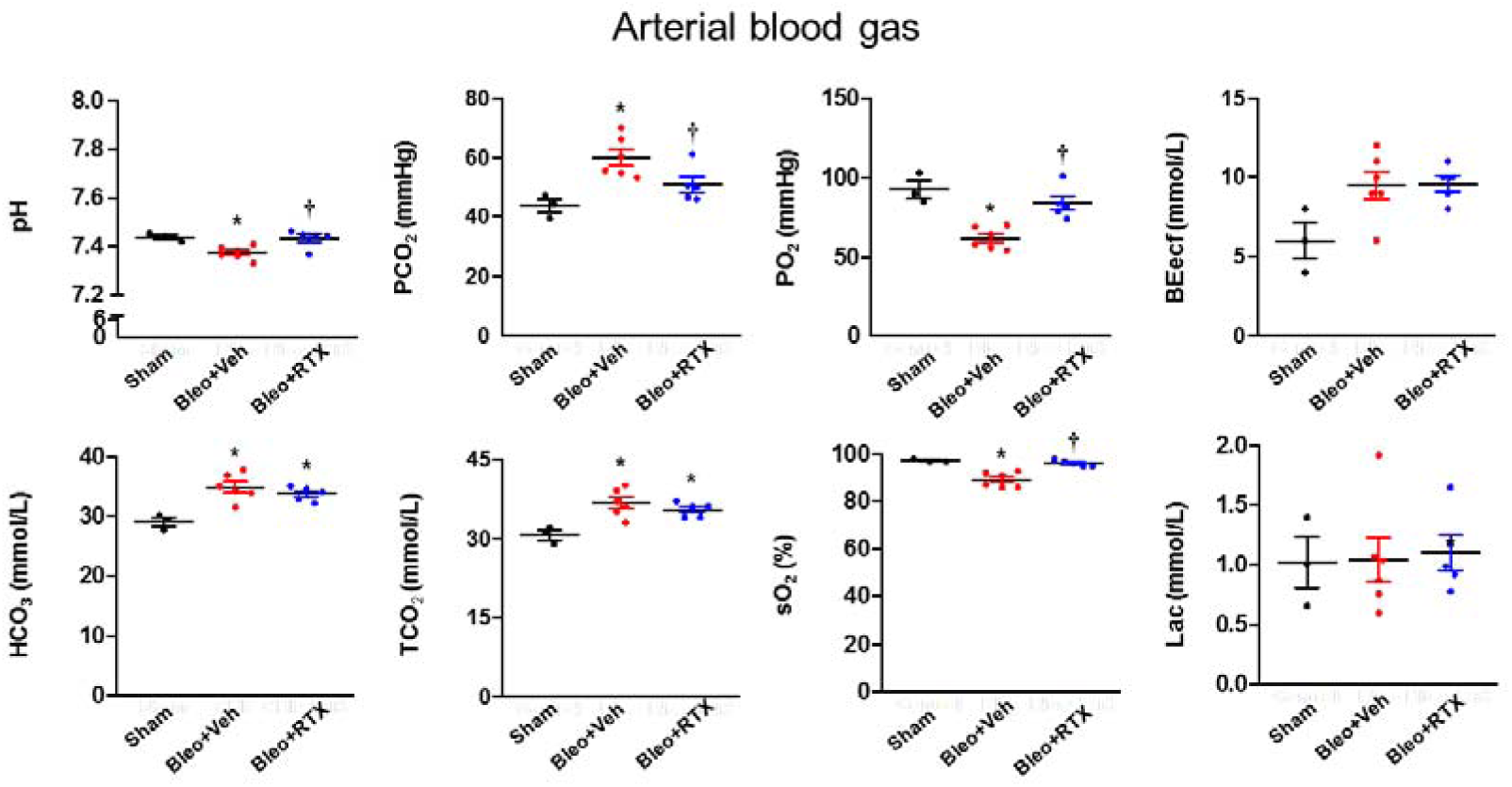
Day 7 arterial blood gases in Sham, Bleo + vehicle and Bleo + RTX rats following intra-stellate administration at day 3 post-injury. PCO_2_, partial pressure of carbon dioxide; PO_2_, partial pressure of oxygen; BE, base excess; TCO_2_, total CO_2_; sO_2_, Oxygen saturation; Lac, lactate. \**P*<0.05 vs. Sham. †, *P*<0.05 vs. Bleo+Veh.

**Figure 7** shows that lung tissue levels of IL-6 and IL1ß were significantly reduced after stellate RTX administration.

**Figure 7.**
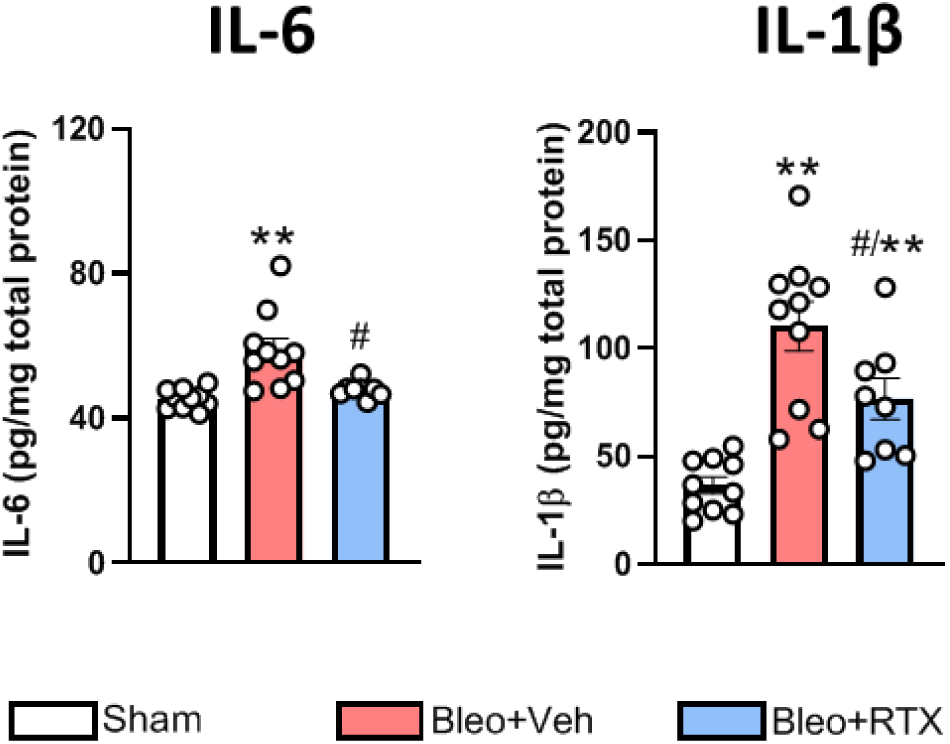
Day 7 lung tissue cytokine levels following day 3 vehicle or stellate RTX administration. IL-6, interleukin 6; IL-1β, interleukin 1 β. **P<0.01 vs. Sham. ^#^P<0.05 vs. Bleo+Veh.

**Figure 8** shows body weight (BW) and individual organ weight among groups. Compared to sham rats, wet lung weight (WLW) as well as the ratio of WLW to BW was significantly higher in the Bleo+Veh rats, which was significantly reduced by intra-stellate injection of RTX. These data suggest that intra-stellate injection of RTX reduces lung edema post Bleo.

**Figure 8.**
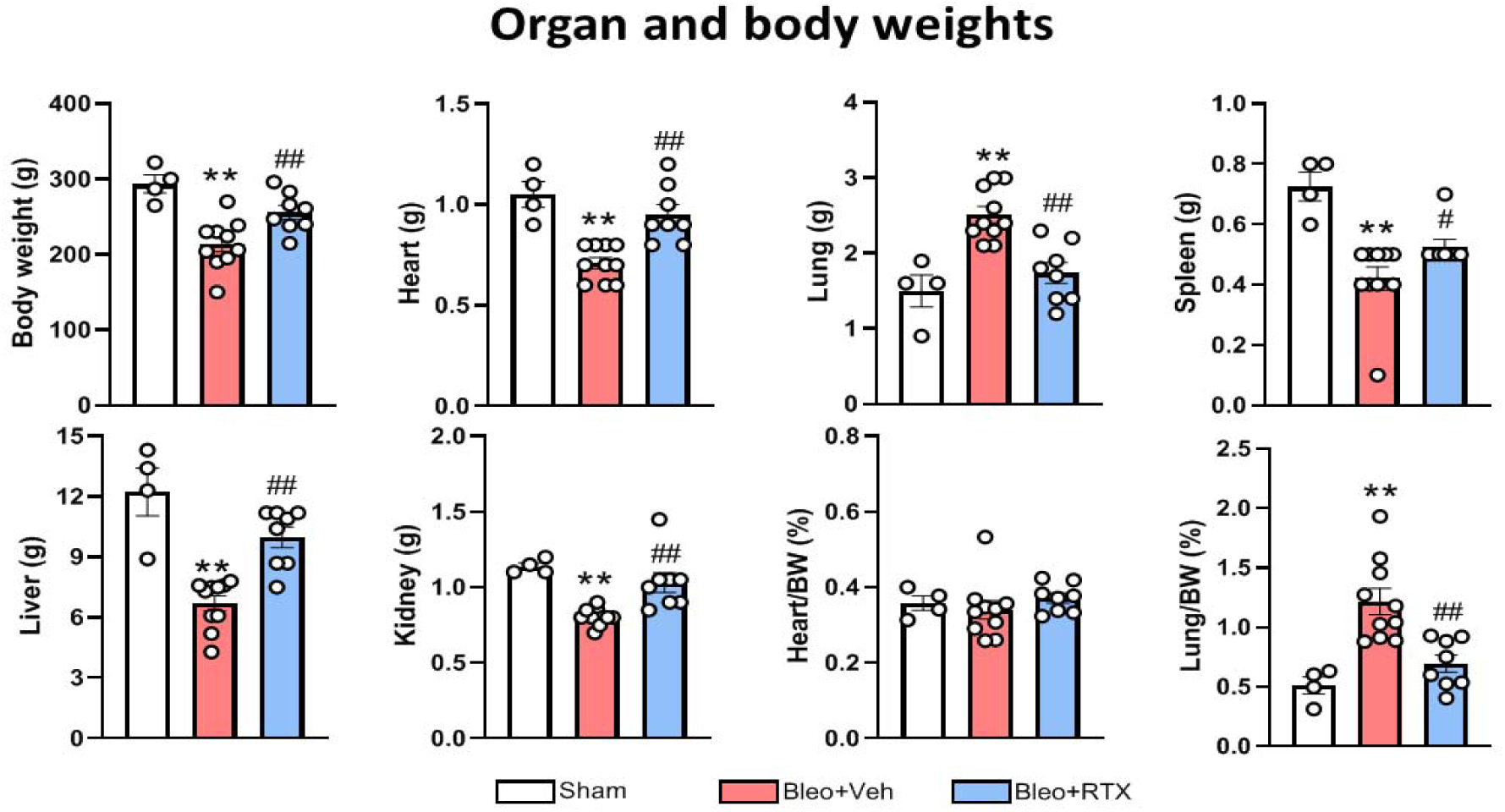
Day 7 body and organs weight in untreated, day 3 vehicle and day 3 stellate RTX treated rats. \*\**P*<0.01 vs. Sham. ^#^*P*<0.05 and ^##^*P*<0.01 vs. Bleo+Veh.

## Discussion

The data from this study provides proof of principle and is highly suggestive of an important role for TRPV1-positive spinal afferent–mediated neuroinflammation in acute lung injury and progression to acute respiratory distress. The evidence provided demonstrates that ablation of TRPV1 sensory afferents in the presence of acute lung injury using RTX delivered by either of two routes that target cardiopulmonary afferents leads to a rapid reduction in lung microvascular permeability and a reduction in tissue and plasma inflammatory markers. While pulmonary function *per se* was not directly measured in this series of experiments, arterial blood gas data strongly suggest an improvement in gas exchange. The improved body weight and reduced lung weight in rats with lung injury after receiving stellate ganglia administration of RTX suggest potential clinical benefits from reduced lung edema, and protective effects for non-pulmonary organs that would otherwise be impacted by the pulmonary triggered systemic inflammatory process.

The lung is innervated by a dual sensory system including vagal and spinal afferents. Both vagal and spinal afferent fibers are composed of A (high conduction velocity) and c-fiber (low conduction velocity) axons. These fibers and their sensory endings express a variety of membrane receptors that mediate ion channel function including traditional Na^+^, K^+^ and Ca^2+^ channels (both voltage and ligand gated). Importantly, non-specific cation channels that are highly permeable to calcium are expressed mostly in small diameter c-fibers.^19, 20^ These include at least 30 members of the Transient Receptor Potential family including Transient Receptor Potential A (TRPA) and TRPV receptors. TRPV1 receptors transduce sensations of heat and neuropathic pain in the periphery. Estimates are that approximately 60 percent of thoracic DRG neurons are positive for TRPV1.^21^ Upon activation, TRPV1 channels are highly permeable to calcium.^19, 20^ High levels of intracellular calcium are toxic and thus damage or kill these specific afferent neurons. Thus, a unique strategy has been developed to modulate the pathological effects of TRPV1 afferent neurons. The ultrapotent neurotoxin, RTX binds avidly to the TRPV1 receptor. After initial stimulation, high intracellular levels of calcium mediate inhibition of neuronal function. Site-specific delivery of RTX can be used to intervene in various conditions to alleviate pain, inflammation, fibrosis and plasma extravasation. It has been shown that RTX-induced TRPV1 sensory afferent deletion can block the afferent-contained neuropeptide release and reduce inflammatory pain.^22^ Cardio-pulmonary spinal afferents can also be targeted with RTX by either application into the epidural space at thoracic levels T1-T4 ^11^ (with some spread to higher and lower segments) or by injection into the stellate ganglia. While DRGs are considered exclusively sensory in nature, the stellates contain soma for sympathetic efferent fibers and fibers of passage for thoracic afferents as they course through DRGs and enter the spinal cord. It should be noted that in humans the stellate ganglia can be easily identified, and that this type of transcutaneous procedure can be performed with fluoroscopic or ultrasound guidance (intra-ganglionic or nerve ‘block’ approach). Compared to epidural delivery that requires relatively larger injection volume (∼80 µl for bilateral injection) to sufficiently cover the T1-T4 DRGs, intra-stellate injection requires a much smaller volume (10 µl for bilateral injection), which reduces the risk of systemic absorption of RTX and allows a higher dose of RTX to be used for local injection.

The current data clearly show that intra-stellate ganglia injection of RTX markedly attenuated lung extravasation post ALI, suggesting that a large proportion of thoracic afferents passing through the stellates innervate the lungs. Taken together, we believe that intra-stellate ganglia delivery of RTX should be a clinically feasible intervention to treat acute lung injury compared to the epidural approach. The preliminary data presented here for epidural administration of RTX provides proof of principle that it reduces plasma extravasation in the lung. The main focus of this study was on the therapeutic effect of the stellate ganglia approach on lung pathology in our ALI rat model.

Prior work from this laboratory has demonstrated that ablation of cardiac TRPV1-positive afferents reduces sympathetic nerve activity and cardiac remodeling in a post myocardial infarction model of chronic heart failure.^23^ TRPV1-expressing cardio-pulmonary afferents participate in a sympatho-excitatory reflex that has been termed the cardiac afferent sympathetic reflex (CSAR)^24^ and the pulmonary afferent sympathetic reflex (PSAR)^25^. The CSAR is augmented in heart failure along with cardiac afferent discharge in response to bradykinin or capsaicin.^23^ Epicardial administration of RTX reduces sympathetic outflow to the heart and kidneys and improves cardiac diastolic function while reducing fibrosis and cytokine content in the heart.^23^ Furthermore, cardiac application of local anesthetic lowers sympathetic nerve activity in anesthetized vagotomized animals suggesting tonic input from these spinal afferents in heart failure.^15, 26^ On the other hand, it has been widely reported that activation of TRPV1-expressing afferents causes secretion of neuropeptides such as substance P (SP) and calcitonin gene-related peptide (CGRP).^27-31^ Released SP, but not CGRP, in sensory endings binds neurokinin (NK) 1 receptors on blood vessels and causes vasodilation and increased vascular permeability that allows loss of proteins and fluid (plasma extravasation) thus promoting the regional accumulation of monocytes and leukocytes contributing to inflammation.^32-34^ In the lung, this process not only impairs alveolar gas exchange but may initiate and exacerbate a fulminant cytokine storm from adjacent cells and from circulating macrophages.^6^ The current study supports the idea that selective ablation of TRPV1 afferents mitigates neuroinflammation in the lung by inhibiting TRPV1 afferent-mediated plasma extravasation, at least in the bleomycin model of ALI. Importantly, although we did not directly measure respiratory parameters such as minute ventilation and respiratory rate in vehicle and RTX treated bleomycin rats, we observed significantly reduced lung weight and improved blood gas parameters including blood pH, pO_2_ and SO_2_ in bleomycin rats treated with intra-stellate ganglia injection of RTX, suggesting an improvement in lung function. Combined with the results shown in the current model of ALI, the potential of RTX to rescue lung function and protect multiple organs from collateral damage due to lung injury triggered inflammation is very encouraging and warrants additional studies as a rescue therapy for patients with lung injuries or infections resulting in inflammatory – mediated pneumonia. Clinical assessments including potential acceleration for return to normal function or long-term protective effects on lung fibrosis should also be explored in upcoming studies.

In the current study, we chose the bleomycin-induced lung injury model to study the role of pulmonary spinal afferent ablation in lung pathology after acute lung injury. Intratracheal injection of bleomycin has been widely used to evoke pulmonary interstitial lesions in animal models.^35, 36^ Bleomycin-induced lung injury is primarily mediated by alveolar epithelial damage resulting in the release of large number of inflammatory cells and cytokines. Following pulmonary insult with bleomycin at day 0, inflammation progresses to peak levels around day three. Pulmonary edema, respiratory distress, body and organ weight loss associated with systemic inflammation is observed up to day 10.^35^ The model was chosen as it closely reproduces important aspects of ARDS including local inflammation, cytokine storm, progression to respiratory distress, and multi-organ impact. The timing of therapeutic intervention (day 3) was also carefully selected to coincide with high levels of inflammatory mediators and lung damage that would be found in infectious disorders including in COVID-19 patients progressing to ventilatory support. We acknowledge a limitation that since the clinical etiology of lung injury is variable (e.g. viral/bacterial infection, chemical and surgical), the bleomycin model does not completely mimic all pathological characteristics of lung injury in humans. While we considered the lipopolysaccharide (LPS)-induced lung injury model, LPS has been shown to have a direct effect on sympathetic and parasympathetic afferent and efferent neurons in addition to its effect on the lung itself,^35, 37-40^ and thus was less suitable for use in this study. As far as a viral infection model is concerned, we have not attempted to directly apply our findings to relevant diseases such as COVID 19, although there may be some phenotypic overlap between the bleomycin lung injury and viral infection models. More work needs to be done to validate the efficacy of RTX in other lung injury models.

## Conclusions

Our data suggest that pulmonary spinal afferent ablation by intra-stellate injection of RTX reduces plasma extravasation and local pulmonary inflammation post bleomycin-induced lung injury which results in improved blood gas exchange. These findings suggests that local stellate application of RTX could be used as a potential clinical intervention to mitigate lung pathology after ALI.

## Sources of Funding

This study was supported by Sorrento Therapeutics Inc. Dr. Hanjun Wang is also supported by Margaret R. Larson Professorship in Anesthesiology. Dr. Irving H. Zucker is supported in part by the Theodore F. Hubbard Professorship for Cardiovascular Research.

## Conflicts of Interest/Disclosures

The lung/RTX project is currently sponsored by Sorrento Therapeutics Inc.

